# MetaPop: A pipeline for *macro*- and *micro*-diversity analyses and visualization of microbial and viral metagenome-derived populations

**DOI:** 10.1101/2020.11.01.363960

**Authors:** Ann C. Gregory, Kenji Gerhardt, Zhi-Ping Zhong, Benjamin Bolduc, Ben Temperton, Konstantinos T. Konstantinidis, Matthew B. Sullivan

## Abstract

**Background:** Microbes and their viruses are hidden engines driving Earth’s ecosystems from the oceans and soils to humans and bioreactors. Though gene marker approaches can now be complemented by genome-resolved studies of inter- (*macro*diversity) and intra- (*micro*diversity) population variation, analytical tools to do so remain scattered or under-developed.

**Results:** Here we introduce MetaPop, an open-source bioinformatic pipeline that provides a single interface to analyze and visualize microbial and viral community metagenomes at both the *macro*- and *micro*-diversity levels. *Macro*diversity estimates include population abundances and α- and β-diversity. *Micro*diversity calculations include identification of single nucleotide polymorphisms, novel codon-constrained linkage of SNPs, nucleotide diversity (π and θ) and selective pressures (pN/pS and Tajima’s D) within and fixation indices (F_ST_) between populations. MetaPop will also identify genes with distinct codon usage. Following rigorous validation, we applied MetaPop to the gut viromes of autistic children that underwent fecal microbiota transfers and their neurotypical peers. The *macro*diversity results confirmed our prior findings for viral populations (microbial shotgun metagenomes were not available), that diversity did not significantly differ between autistic and neurotypical children. However, by also quantifying *micro*diversity, MetaPop revealed lower average viral nucleotide diversity (π) in autistic children. Analysis of the percentage of genomes detected under positive selection was also lower among autistic children, suggesting that higher viral π in neurotypical children may be beneficial because it allows populations to better ‘bet hedge’ in changing environments. Further, comparisons of *micro*diversity pre- and post-FMT in the autistic children revealed that the delivery FMT method (oral versus rectal) may influence viral activity and engraftment of *microdiverse* viral populations, with children who received their FMT rectally having higher *microdiversity* post-FMT. Overall, these results show that analyses at the *macro-level* alone can miss important biological differences.

**Conclusions:** These findings suggest that standardized population and genetic variation analyses will be invaluable for maximizing biological inference, and MetaPop provides a convenient tools package to explore the dual impact of *macro*- and *micro*-diversity across microbial communities.

## Introduction

Microbiology has experienced a revolution as sequencing and computational advances have enabled the cultivation-independent study of microbial and viral communities across diverse ecosystems. These studies have revealed the importance of the ‘microbiome’ and its viruses as critical drivers whose metabolisms and impacts alter nutrient, metabolite and energy flows that dictate human health and ecosystem outputs (e.g., Falkowski *et al*. 2008, Kinross *et al*. 2011, Knight et al. 2017). Pragmatically, the sequence space exploration has helped rewrite foundational taxonomic rules even to the point of genomes alone being sufficient (Roux *et al*. 2018, Parks *et al*. 2020). Though early studies relied upon gene marker derived amplicons and could answer “who is there” questions (e.g., Lane *et al*. 1985, Woese 1994), sequencing and analytical advances have led to increasingly improved assembles such that genome-resolved, population-level analyses now get beyond “who is there” to understand metabolism and even mechanism (e.g., Koh *et al*. 2016, Roager *et al*. 2016, Woodcroft *et al*. 2018, Solden *et al*. 2018, Martinez-Guryn *et al*. 2018). This transformation has happened rapidly, with catalogs of tens to hundreds of thousands of microbial and viral metagenome-assembled-genomes now emerging across diverse environments (e.g., Parks *et al*. 2017, Delmont *et al*. 2018, Pasolli *et al*. 2019, Almeida *et al*. 2019, Nayfach *et al*. 2019, Emerson *et al*. 2018, Shkoporov *et al*. 2019, Gregory *et al*. 2019, Gregory *et al.,* 2020). Beyond such inter-population (*macro*-diversity) community questions, recent advances are now also providing a window into intra-population (*micro*diversity) variation. These latter observations provide complementary information by establishing niche-defining gene sets, as well as how genetic drift and selection shape populations and communities (e.g. Schloissnig *et al*. 2013, Gregory *et al*. 2019, García-García *et al*. 2019).

A major challenge when assembling fragmented DNA from complex communities is assembling short-reads into biologically meaningful ‘genomes’ that represent ecologically and evolutionarily relevant populations. At this point, however, there are several improved population definitions that account for ecological and evolutionary theory (Cordero & Polz 2014) and have been extrapolated and assessed to varying degrees community-wide (Caro-Quintero & Konstantinidis 2012, Gregory *et al*. 2016, Bobay & Ochman *et al*. 2018, Olm *et al*. 2020). Many of the remaining criticisms, e.g., chimeric ‘franken-genomes’, are being increasingly addressed by the rapidly advancing capabilities enabled by long-read sequencing and hybrid assembly approaches (e.g. Warwick-Dugdale *et al*. 2019, Moss *et al*. 2020). Thus, researchers studying microbes and viruses in complex communities have or soon will have datasets that are ready for genome-resolved population-based studies where high-fidelity assemblies and base calls can be expected.

Once assembled, several obstacles remain to establish intra-population biological inferences. *First,* population genetic methods rely on defined genotypes of individuals within a population with equal coverage across each base within a sequence. These are conditions not satisfied in metagenomes as their populations have unequal coverage and are assembled from many individuals within a population. To date, researchers have developed many methods to overcome these issues. The most common ones try to resolve each individual’s genotype within the population by linking single nucleotide polymorphisms (SNPs) into strain-level genotypes (Ahn et al 2014; Hong et al 2014; Sankar et al 2016; Albanese et al 2017; Tamburini et al 2018, Luo et al 2015; Sahl et al 2015, Nayfach et al 2016; Quince et al 2017; Fischer et al 2017; Smillie et al 2018) or use strain proxies (Eren *et al*. 2015, Scholz et al 2016, Greenblum *et al*. 2015, Eren *et al*. 2015), but these are difficult to apply to or are insufficient for community-scale studies across bacteria and viruses. A *second* obstacle is that analyzing both *macro*- and *micro*diversity in these modern datasets have scaling and standardization issues, and user-friendly bioinformatic tools are not yet available. For the latter, while several bioinformatic tools have emerged, they require intensive data manipulation prior to use and few do more than one type of analysis (Eren *et al*. 2015, Nayfach *et al*. 2016, Costea *et al*. 2017, Olm *et al*. 2020, see **Table 1** in **Implementation**). This creates a research barrier for microbiologists that are light on computational skills, which could be alleviated by a tool that provided a single interface to analyze and visualize macro- and micro-diversity patterns in metagenomic data.

To fill this gap, we introduce the multi-functional bioinformatic pipeline ***MetaPop***, written in R and bundled in the bioconda package ‘metapop’, that analyzes and visualizes microbial and viral community metagenomic sequence data at both the inter- (macrodiversity) and intra- (microdiversity) population levels. MetaPop can be easily utilized by beginner microbiologists with little training. In this sense, the pipeline complements existing bioinformatic pipelines, such as Anvi’o (Eren *et al*. 2015) and metaSNV (Costea *et al*. 2017), which require users to modify the code, train, consult detailed tutorials, and continually provide input while running the pipeline. MetaPop’s distinctive features include: (1) it combines both *macro*- and *micro*-diversity analyses into a single easy-to-use pipeline, (2) all of MetaPop’s functions and parameters are called in a single command-line that processes and analyzes the input data from start to finish (with the option to run steps independently), and (3) it improves adaptive selection (pN/pS) results by determining if SNPs are linked at the codon level. MetaPop is fully documented and maintained by the developers at https://github.com/metaGmetapop/metapop.

## Implementation

### Technical overview of how MetaPop works: input, data processing, and output

MetaPop has three inputs: (1) a genome FASTA file, (2) a tab-delimited file of the number of reads or base pairs per metagenomic library, and (3) one BAM file per metagenomic library of read alignments (mappings) to the reference genomes. MetaPop is best applied when each BAM file is derived using reads from a single metagenomic community, rather than reads pooled from multiple communities, in order to prevent the formation of hybrid populations that could violate underlying assumptions of population genetic inferences. With these inputs, MetaPop analyzes the data in three steps (**Figure 1**):

1. Pre-processing
2. *Macro*diversity and codon bias analyses
3. *Micro*diversity analyses

**Figure 1.**
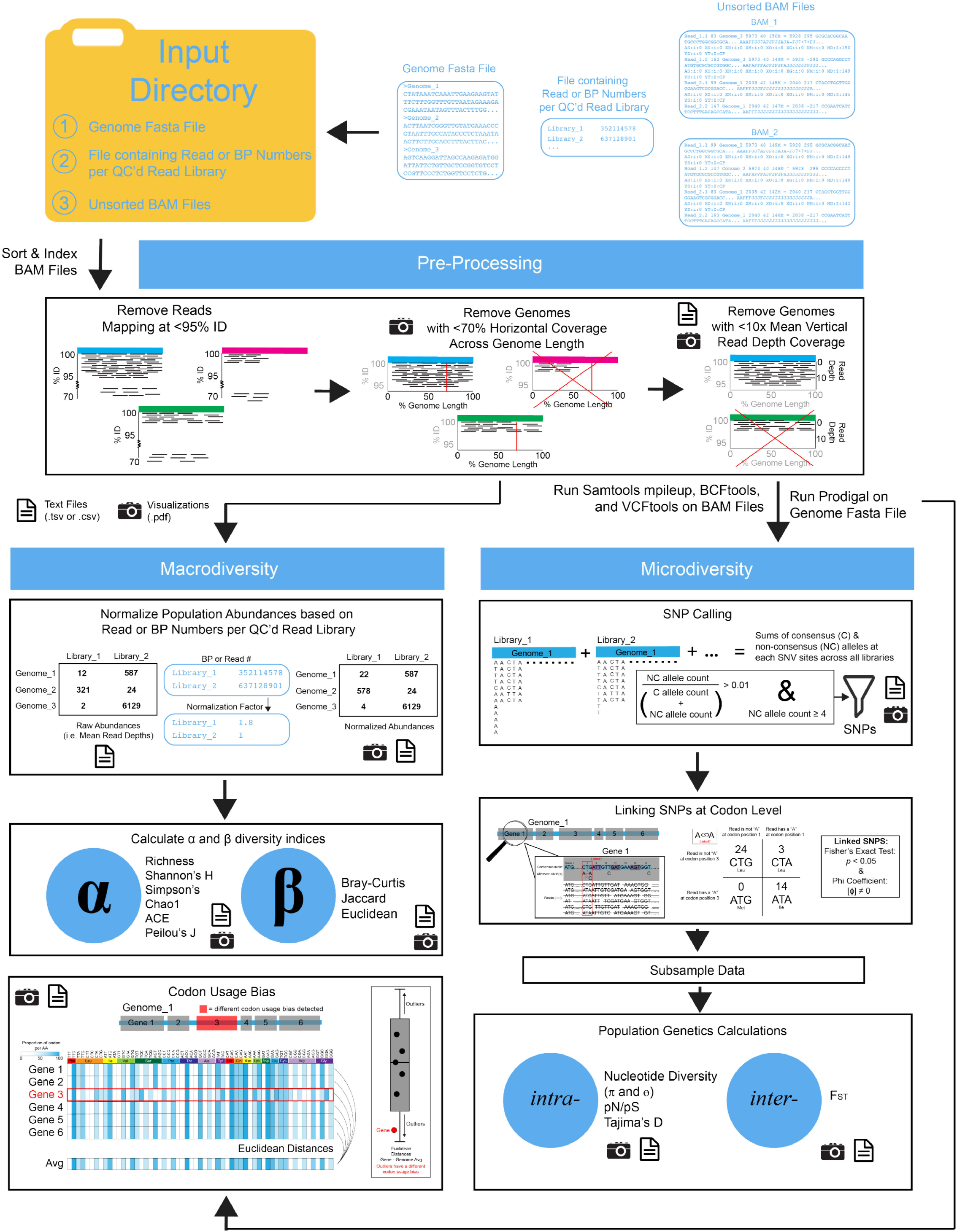
MetaPop Pipeline overview. Metapop requires three primary inputs (a genome fasta file, file with the number of reads or bps per library, and unsorted BAM files). The BAM files are sorted and indexed and preprocessed. The output of preprocessing goes through the macrodiversity or microdiversity arms of the pipelines. Codon usage bias is calculated as well and can be calculated independent of the whole MetaPop pipeline.

To clearly and explicitly lay out the capabilities of MetaPop compared to existing complementary bioinformatic pipelines (Eren *et al*. 2015, Nayfach *et al*. 2016, Costea *et al*. 2017, Olm *et al*. 2020), we provide a summary **(Table 1)**. With the exception of the filtered reads, which are in binary alignment format, data outputs are tab-delimited files and visual outputs vectorized images stored in PDFs. The final tab-delimited files include: (*i*) percent alignment and read length for every read in each sample, (*ii*) percent of positions covered by reads and average depth of coverage for each genome, per BAM file, (*iii*) the raw and normalized genome abundances, calculated *(iv)* α- and (*v*) β- diversity values, *(vi)* genes with different codon biases, (*vii*) single nucleotide variant (SNV) calls and pileup files over SNV positions, (*viii*) called single nucleotide polymorphisms (SNPs), split into those which appear on genes and those which appear in intergenic regions, (*x*) linked SNP results, and (*xi*) *intra*- and (*xii*) *inter*- population genetic calculations. The visualization output include: (*i*) overall summaries of read filtering, (*ii*) summary plots showing BAM file genome coverage and depth statistics, (*iii*) a heatmap of normalized genome abundances, (*iv*) scatterplots of α-diversity values, (*v*) ordination plots of β-diversity results, (*vi*) bar plots of codon position of detected SNPs, (*vii*) visualization of nucleotide diversity and codon bias per each gene for each genome, and identification of positively selected genes, and (*viii*) heatmaps of FST per genome across samples.

### Step 1: Pre-processing

Though user-customizable, MetaPop defaults to a 95% nucleotide identity (ID) cut-off, but can be changed by the user, to define population boundaries, guided by studies exploring sequence space boundaries between microbial and dsDNA viral populations (Konstantinidis *et al*. 2006, Caro-Quintero & Konstantinidis 2012, Gregory *et al*. 2016, Bobay & Ochman 2017, Gregory *et al*. 2019, Roux *et al*. 2019, Olm *et al*. 2020). During pre-processing, input BAM files are first sorted and indexed and reads that map at <95% ID to a reference genome or which are shorter than 30 base pairs are removed. Genomes that pass either a length and/or percentage coverage minimum after this initial filtering are considered present or ‘detected’ in a given sample and move on to *macro*diversity analyses (Couto *et al*. 2018, Gregory & Zayed *et al*. 2019). If the user flags the genome fasta dataset as complete microbial genomes (- complete_bact), MetaPop will use a default detection cut-off of ≥20% genome length covered to consider the genome in further analyses (Castro *et al*. 2018, Couto *et al*. 2018). If the user flags the genome fasta dataset as viral (-viral) or fragmented microbial contigs (-frag_bact), Metapop’s default detection cut-offs require at least ≥5kbp genome length covered in genomes >5kbp and ≥70% length for genomes <5kbp (Gregory & Zayed *et al*. 2019). All length cut-offs, however, can be adjusted by the user.

Once a population is detected by these above criteria, MetaPop calculates its relative abundance based on mean nucleotide coverage across the genome (see ***Macrodiversity Analyses*** section for more details). Loci with coverages below the 10th and above the 90th percentile are excluded from this assessment to prevent skewing of abundances from fast evolving regions, such as genomic islands, and spurious recruitment of reads to highly conserved regions, respectively (Kunin *et al*. 2008). Importantly, users can customize any of these cut-offs for percent identity to define populations, horizontal coverage of the genome to ‘detect’, and the quantiles for minimizing abundance data skew.

### Step 2: Macrodiversity and Codon Bias Analyses

#### Data processing and calculating alpha and beta diversity indices

Macrodiversity is the measure of population diversity within a community. While some diversity measurements rely strictly on the presence or absence of populations (such as richness and Jaccard distances), many rely on the relative abundances of populations between communities (such as Shannon’s H, Simpson’s, and Bray-Curtis distances). Importantly, metrics that rely on relative abundances have been shown to be more robust for metagenomic data because they are less susceptible to uneven sampling of rare taxa (Haegeman *et al*. 2013). Thus, the raw abundances calculated during the pre-processing step must be transformed in order to allow for differential abundance testing. MetaPop proportionally normalizes per-sample abundances to those for the library with the highest number of either the number of reads or base pairs (selected by the user). For example, if library A has 1.5 million reads and library B has 2 million reads, all the raw population abundances in library A are multiplied by 1.33 to in order to proportionally scale the abundances to the library with the highest number of reads. If more than one sequencing technology was used to create the different metagenomes and this resulted in vastly different read lengths, we recommend using base pair counts for the normalization step. Normalized genome abundances for each metagenomic sample are used to calculate macrodiversity measurements with the “vegan” R package. If the input consists of a single BAM file, α-diversity *(within* community) indices – richness, Chao1, ACE, Shannon’s H, Simpsons, inverse Simpsons, Fisher, and Pielou’s J – for that community are calculated. With multiple BAM input files, β-diversity (*between* community) indices –Jaccard, Bray-Curtis, and centered log-ratio-transformed Euclidean distances – between all communities are also calculated. Importantly, MetaPop also outputs the raw abundances, so that the user can normalize their own data.

#### Codon Bias Analyses

Microbial and viral populations often have distinct codon biases for translational optimization (Ikemura 1985). Genes with codon usages different from the rest of the genome often have been recently horizontally transferred (Lawrence & Ochman 1998), have different temporal regulations (Shin *et al*. 2015), or are highly expressed (Sharp & Li 1987). MetaPop predicts putative genes using Prodigal (Hyatt *et al*. 2010) for all genomes in the reference genome FASTA file and identifies the codon usage for each gene within a genome and then calculates the codon bias for each gene. The bias for each codon per amino acid across every gene in the genome is then averaged to create the average codon bias. Each gene’s codon usage is compared using Euclidean distances to the average codon bias. Genes with Euclidean distances greater than 1.5 times the interquartile range, the standard constant for discerning outliers (reviewed in Whitley & Ball 2001), of Euclidean distance for that genome are considered potential outliers and are marked as having aberrant codon usage for their respective genome.

### Step 3: Microdiversity Analyses

#### Data Transformation

Microdiversity is the measure of genetic diversity within a population. In natural communities, where populations are represented at different abundance levels, only genomes with enough data can be evaluated. Thus, by default, genomes with <70% length of their genome covered and <10x average read depth coverage are excluded from these analyses to ensure that there is enough coverage to accurately call SNPs and to assess contig-level microdiversity. The 10x value was selected because prior work revealed that downsampling read depth to 10x did not statistically significantly impact downstream microdiversity calculations (Schloissnig *et al*. 2013). While deeper sequencing is now resulting in high coverage for many microbial and viral populations, it is also uncovering rare low-abundance species that remain with low coverage (Knight *et al*. 2012, Gregory *et al*. 2019). Thus, in order to compare these low and high coverage species, downsampling remains important. Users can set this parameter if they want to be more stringent or relaxed in the number of populations that pass to the microdiversity analyses step. Prior to SNP calling, MetaPop identifies SNVs within each population per BAM using the mpileup tool in samtools (Li *et al*. 2009) and BCFtools (Danecek *et al*. 2017) in order to obtain per-position variant information, followed by removal of low (PHRED < 20 by default) variant quality score calls. Importantly, decreasing or increasing the PHRED threshold for variant quality scores increases or decreases, respectively, the number of SNPs called and the downstream nucleotide diversity values. SNPs are identified using two methods, either a (1) global- or (2) local-approach. For global SNPs calls, the base pair coverage for each SNV position per genome is pooled across all metagenomes and the consensus allele verified. Alternate alleles that make up ≥1% of the base pair coverage for that position (1000 Genomes Project Consortium 2010) and represented by at least 4 reads are considered true SNPs (Schloissnig *et al*. 2013). For local SNP calls, the set of true positions identified in the global calls are reduced to the set of SNV positions identified in each BAM individually. SNV sites only observed in other BAM files are ignored.

Identified SNPs are cross-referenced with gene calls and assigned as either genic or non-genic. If genic, their position within each codon per gene is determined. Due to redundancy at the third position in codons that allows multiple codons to code for the same amino acid, most true SNPs should be at the 3^rd^ position of the codon. MetaPop outputs all of the SNPs called and their codon positions if genic. MetaPop will issue a warning if there are more SNPs in the 1^st^ and 2^nd^ positions of a codon. Lastly, the global verified consensus allele per each SNP position is replaced as the consensus allele in each reference genome.

#### SNP linkages in codon variants and downsampling

SNPs are tested for local linkage at the codon level to identify codon variants by evaluating their co-presence within reads in each BAM file. Multiple programs try to link SNPs across the genome into strain haplotypes using the reads (Nayfach et al 2016; Quince et al 2017; Fischer et al 2017; Smillie et al 2018). However, given that shotgun sequencing read lengths are shorter than most gene and genome lengths, it makes it difficult to resolve genotype patterns that span across more than single read length. Assessing linkage across small sequences that can be contained with a single read, nonetheless, provides the strongest evidence of linkage. MetaPop tests for linkage at the codon level due to its importance for studying protein evolution. The linkage of SNPs, for example, at positions 1 and 3 within a codon can code for a completely different amino acid then if each SNP independently arose. Further, codons are short enough to be contained with a single read, which allows us to accurately test the linkage between or independence of SNPs shared within a codon. The resulting codon variants that code for different amino acids, most often those that have two SNPs or SNPs in the 1st or 2nd position of the codon, and their resulting impact on protein structure have recently become an active area of research in metagenomes (Delmont *et al*. 2019). Recent work, however, identified these codon variants by filtering for highly abundant codon variants already contained within 20 reads (Delmont *et al*. 2019). To the best of our knowledge, MetaPop is the first program to try to statistically link SNPs at the nucleotide level into these codon variants.

In order for MetaPop to link the SNPs at the codon level, SNPs that localize in the same codon on the same gene are selected as candidates for linked SNP identification. The original reads covering the positions of the candidates from their respective genomes are collected from each BAM file, and the codons relevant to the candidates are extracted from those reads. SNPs within the same codon are tested in pairs. If more than two SNPs occur within a codon, pairwise tests are performed between each combination of pairs. A contingency table of the frequencies of the extracted codons with both SNPs, the number of extracted codons with one SNP, the number of extracted codons with the other SNP, and those containing no SNPs is produced. Fisher’s exact tests and phi coefficients are calculated. The linked SNP candidates are classified as either “linked” (Fisher’s *p*-value < 0.05, φ> 0) meaning the SNPs occur together as a set disproportionately, “independent” (Fisher’s *p*-value < 0.05, φ < 0), meaning the presence of one of the candidates excludes some of all of the others in that set disproportionately, or as “ambiguous”, meaning that they occur together or separately at apparent random, or that there is insufficient data to classify them otherwise. In ambiguous cases, SNPs are treated as independent for downstream analyses.

SNP frequencies are subsampled down to 10x coverage proportionate to the frequency of different SNPs per site while maintaining SNPs linkages. This stage normalizes the probability of a variant occurrence by chance across variant sites within a population genome. It also rarefies all the SNP frequencies across all genomes and samples allowing for differential SNP frequency testing. While the user can adjust the subsampling level, we recommend that you subsample down to the same average read depth cut-off.

#### *Population Genetic Calculations:* θ, π, F_ST_, pN/pS, and Tajima’s D

The subsampled SNPs are then used to assess population level genetic diversity and explore protein and genome evolution. This will occur with both global and local SNP calls. If the population occurs in only one BAM file, only *intra*-population diversity (within population *micro*diversity) – expected nucleotide diversity (θ; Watterson 1975) and the observed nucleotide diversity (π; Nei & Li 1979) – are calculated. Both θ and π are calculated at the individual gene and whole-genome levels. Because we use a default minimum genome coverage of 70%, not every SNP position for a population will be covered within a BAM file. To correct for this, we use the following equation to estimate θ:

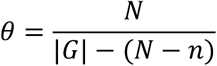

where *N* is the total number SNP positions within a gene or genome, *n* is the number SNPs covered within a metagenome, and *|G|* is the total gene or genome length. To estimate π, we modified the Schloissnig et al (2013) equation

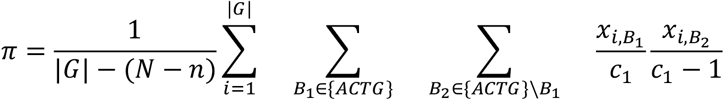

where *N* is the total number SNP positions within a gene or genome, *n* is the number SNPs covered within a metagenome, |*G*| is the total gene or genome length, *x_i_,B_j_* is the number of nucleotide *B_j_* seen at position *i* and *c_i_* the coverage at position *i* in the gene or genome. If the population occurs in more than one BAM, F_ST_ (Wright 1965; between population *micro*diversity) is calculated. Because F_ST_ requires comparing the nucleotide diversity per site across two metagenomes, we chose to keep the total genome length as the common denominator given that the SNP coverage may vary between both metagenomes. The implemented equation for F_ST_ is directly from Schloissnig et al 2013.

To explore selective pressures on specific genes, MetaPop uses two methods: pN/pS (Schloissnig et al 2013) and Tajima’s D (Tajima 1989). The implemented equation pN/pS is directly from Schloissnig et al 2013 except it factors in codon-constrained SNP linkages. Tajima’s D is calculated using the original equation (Tajima 1989), but using the π value calculated above, the number of SNP positions within a gene as the number of segregating sites, and the ceiling mean read depth for the number of sequences.

## Results & Discussion

### Biological Evaluation of MetaPop

In order to test MetaPop, we ran the pipeline on three previously published datasets, a synthetic dataset representing mock bacterial communities and two biological virome dataset with natural variation (i.e., beyond that in the mock community). The synthetic dataset is composed of 30 mock, bacterial metagenomic communities of different known-proportions of *Staphylococcus aureus, Staphylococcus epidermidis,* and *Bacillus subtilis* strains (Sankar *et al*. 2016; see **Table S1**). The first biological virome dataset included 131 of the viromes from the Global Ocean Viromes 2 (GOV2; Gregory *et al*. 2019) dataset. This dataset was the first dataset to assess *micro*diversity in metagenome-assembled viral genomes at community-wide scale and provided the methodological backbone of MetaPop. The second biological virome dataset was composed of gut viromes from 12 autistic children that underwent fecal microbiota transfers and 6 neurotypical children (Kang *et al*. 2017; see **Table S2**). The default visualization outputs of MetaPop for the second biological virome dataset can be seen in **Fig. 2**.

**Figure 2.**
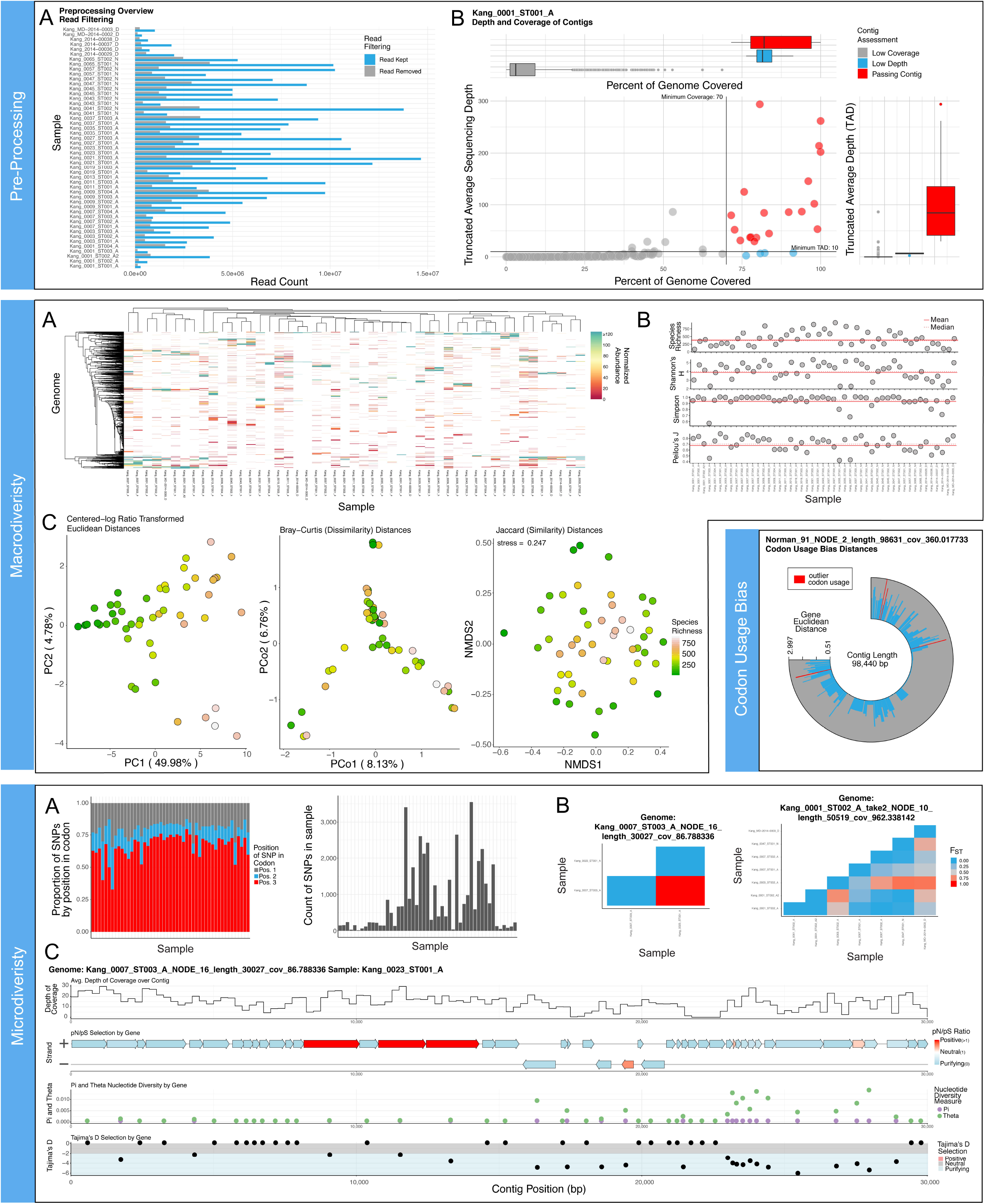
MetaPop Visualization Outputs from our autism biological virome dataset. **Preprocessing** visualizations include **(a)** bar plot showing how many reads were kept and removed following removal of reads below a 95% ID cut-off across all samples, **(b)** scatter and bar plot composites (1 example shown) reported per sample showing how many genomes pass the horizontal and vertical coverage cut-offs, and **(b-inset)** donut plots summarizing the total number of genomes passing the different horizontal and vertical coverage cut-offs per sample. **Macrodiversity** visualizations include **(a)** heatmap summarizing the normalized abundances of covered genomes across the different samples (the max value on the color scale reported is the 75% quantile of all abundances to allow low abundance genomes to be better displayed; another heatmap not shown is also created showing full range of abundances), **(b)** scatter plots per each alpha diversity index (4 examples shown) showing the alpha diversity value across all samples with horizontal lines showing the mean and median values, **(c)** ordination plots (PCA, PCoA, and NMDS) of all centered-log ratio transformed Euclidean distances, Bray-Curtis distances, and Jaccard distances, respectively (all distances are plotted using the 3 ordination methods by default). The color of each circle represents the species richness within each sample. The **codon usage bias** visualization is a circular bar plot per genome (1 example shown) showing the Euclidean distance of each gene from the average gene codon bias. Gene with outlier codon biases are displayed in red. **Microdiversity** visualizations include: **(a)** Stacked bar plot (right) and standard bar plot (left) showing the distribution of SNPs across codon position and total number of SNPs per sample. **(b)** F_ST_ heatmaps per genome (2 examples shown) showing the population differentiation per genome across all samples it has coverage within. **(c)** Genome plot composites for each genome in each sample where it has coverage (1 example shown) with four different tracks from top to bottom showing a line graph of the depth coverage of the genome, a genome plot of the genome with coloration of genes showing pN/pS results, a scatter plot showing π and θ values, and lastly a scatter plot showing Tajima’s D values with the color background showing whether the value is indicative of selection.

#### MetaPop reproduces macrodiversity patterns in silico mock communities

We first tested MetaPop’s default settings to accurately determine community composition and to calculate *macro*diversity values across the 30 mock bacterial metagenomic communities. The communities are of varying known proportions of three distinct strains of *S. aureus,* three distinct strains of *S. epidermidis,* and a single strain of *B. subtilis* (Sankar *et al*. 2016) and have varying numbers of reads, from ~2 million reads (communities 1-10), ~3 million reads (communities 11-20), ~6 million reads (communities 21-30). This dataset is practical to test *macro*diversity calculations in MetaPop because the community is composed of two dominant closely related bacterial species-level populations (*S. aureus* and *S. epidermidis*) that share >80% ANI and a more distantly related, rare, species-level population (*B. subtilis)* (**Fig. S1**). This taxonomic combination and different simulated sequencing depth enabled us to determine whether MetaPop could distinguish between closely related populations, and whether increased *micro*diversity within a population as well as sequencing depth impacted our ability to correctly assess *macro*diversity.

Across the 30 communities, bacterial population relative abundances were almost identical to the simulated proportions with only a 0.98- and 1.08- mean fold change differences of the *S. aureus* and *S. epidermidis* species, respectively, and a 0.26-fold change average difference of the *B. subtilis* which is simulated to represent a rare taxa across the communities (**Fig. S2A**). This fold-change difference is similar to or less than known quantitative biases, such as 10% divergence in alpha- and beta-diversity values seen in other metagenomic analyses for viruses (Roux *et al*. 2018) and ~3-9% divergence the microbes if genome length is accounted for (Nayfach & Pollard 2015). The number of strains within each population and the number of reads did not impact detection of different populations (**Fig. S2A**). MetaPop estimates of α- diversity (**Fig. S2B;** all α-diversity indices: Wilcoxon *p* > 0.05) and β-diversity (Bray-Curtis dissimilarity) did not significantly differ from the actual values (**Fig. S2C;** Mantel’s test *p* > 0.05). Thus, despite the minor fold-change differences in community composition and difficulty in accurately detecting the abundances of rare taxa, MetaPop is generally able to accurately assess the community composition and *macro*diversity biological trends.

#### MetaPop’s codon usage bias analyses detects highly expressed and horizontally transferred genes in Staphylococcus aureus

MetaPop also looks for variation in codon biases among genes within each genome. Genes with different codon usage are often associated with horizontal gene transfer, high expression, or different temporal regulation of expression (Lawrence & Ochman 1998, Sharp & Li 1987, Shin *et al*. 2015). To determine the biological validity of MetaPop’s codon’s usage analysis, we evaluated codon bias across all 7 strains in the mock community. We choose to focus our analyses on the genome of an ST5 methicillin-resistant strain of *S. aureus* (see full list of codon bias outliers in **Table S3**) because it is a well-studied human pathogen with known regions of the genome that were horizontally transferred (Malachowa *et al*. 2010, Alibayov *et al*. 2014) or highly expressed (Karlin *et al*. 2001, Malachowa *et al*. 2011, Peschel & Otto 2013). Importantly, MetaPop, in its default settings, is conservative (but user changeable) because it compares each gene’s codon bias to the average codon bias across the whole genome and will, thus, underestimate the number of genes with different codon usage. Given this conservative approach, we could only validate how many genes with a detected difference in codon bias were either mobile elements or highly expressed genes. Across the *S. aureus* ST5 strain, the vast majority (71%; 104 out 149) of the genes detected to have outlier codon usage have no known function. Of the remaining 45 annotated genes MetaPop detected with outlier codon usages, 20% are genes on known mobile elements (so prone to HGT) or are thought to be horizontally transferred and 47% are known to be highly expressed (**Fig. S2D**). The identified known mobile elements include many toxin-antitoxin system genes (Malachowa *et al*. 2010, Alibayov *et al*. 2014) and putatively transferred DltX and DltC proteins involved in wall teichoic acid, as well as poly(glycerol-phosphate) alpha-glucosyltransferase, a type of glycosyltransferase (Li *et al*. 2015). The highly expressed genes include ribosomal proteins (Karlin *et al*. 2001), genes involved in transcription and translation such as elongation factor Tu (Karlin *et al*. 2001; Soufo *et al*. 2010), chaperones (Karlin *et al*. 2001; Bae *et al*. 2000; Duval *et al*. 2010), and all the phenol-soluble modulins (PSMα1-4 and PSMβ; Peschel & Otto 2013). Thus, MetaPop, in its default conservative settings, will not identify all horizontally transferred genes or highly expressed genes, but it provides an important first look at potential targets for further study.

#### MetaPop reproduces microdiversity patterns in the global oceans virome 2 (GOV2) dataset

Using the GOV2 dataset, we next evaluated MetaPop’s ability to assess *micro*diversity values and trends (**Fig. S3**). The original GOV2 paper explored *micro*diversity in the form of average nucleotide diversity (π) per sample by randomly subsampling the π values of different viral populations in each sample and averaging those values. We replicated these methods with the π values calculated using MetaPop where SNPs were called locally when they had a differential base with a PHRED score ≥30 (see Methods). As a result, we ran MetaPop using PHRED≥30 and its default of ≥20 on the GOV2 dataset.

Importantly, MetaPop calculates π slightly differently than the method used in the original analyses of the GOV2 dataset. The original method used the exact equation derived from Schloissnig *et al*. 2012 which divides the calculated nucleotide diversity by the total genome length to obtain π. Because of unequal coverage across genomes in each sample, SNP positions are often not covered so it is impossible to assess the diversity at that site. As a result, MetaPop subtracts the number of SNP positions not covered from the genome length prior to dividing the nucleotide diversity in order to calculate π. Thus, π values from MetaPop will be slightly higher than values using the Schloissnig *et al*. 2012 equation. As expected, MetaPop’s average π using the same SNP calling thresholds (PHRED≥30 and local SNP calls) were slightly higher (median 1.33 fold-change) than the original GOV2 average π (**Fig. S3A, left**). Due to the differences in random subsampling, there were also clear deviations between the original GOV2 average π values and the MetaPop derived values. Nonetheless, the original GOV2 average π and MetaPop’s average π still strongly correlated (**Fig. S3A, right;** linear regression: R^2^ > 0.62), indicating that despite higher average π and slight fluctuations in average π derived from the random subsampling process, the biological *micro*diversity patterns are still being captured.

The SNP calling approach (global versus local) and PHRED score (i.e. a measure of the quality of the called nucleotide) can also impact downstream π values. Global SNP calling, for example, incorporates all SNP loci that were identified in any sample in the dataset into the π calculation for each sample (even if it was not called as an SNV for that exact sample), which will increase π. Using the GOV2 dataset, we see just that with PHRED≥30 global SNP derived average π having a median 4.32-fold increase from the original GOV2 average π values calculated using PHRED≥30 and local SNP calls (**Fig. S3B, right**) and a median 3.19- fold increase over MetaPops’s average π using PHRED≥30 and local SNP calls (**Fig. S3E, right**). Further, decreasing the minimum PHRED score requirements allows more potential SNVs and thus SNPs to be called per sample and, thus, the average π values should be higher. As expected, we see that using a PHRED≥20 global SNP call approach increases average π values by a median 5.88-fold increase from the original GOV2 average π values calculated using PHRED≥30 and local SNP calls (**Fig. S3C, left**) and 1.32 fold increase from MetaPop’s PHRED≥30 global SNP approach (**Fig. S3D, left**). Importantly, regardless of SNP calling approach or PHRED score cut-off, the π calculated using the original GOV2 approach or MetaPop’s approaches are all strongly correlated (**Fig. S3A-E, right;** linear regression: R^2^ > 0.48 to 0.68). Further, analyses of larger *micro*diversity trends across ecological zones in the ocean defined in the original GOV2 analyses were also able to be replicated using both PHRED score cut-offs testing and a global and local approach (**Fig. S3F**). Taken together, MetaPop is generally able to accurately derive *micro*diversity values and biological trends.

#### MetaPop’s condon-constrained linkage of SNPs improves detected of positively selected genes

Using the two biological datasets, we evaluated the impact of MetaPop’s novel codon-constrained SNP linkages on pN/pS selection analyses. The original pN/pS equation (Schloissing *et al*. 2012) calculates the number of non-synonymous and synonymous codons without first evaluating if SNPs within the same codon are linked. If the two codon-constrained SNPs are linked, with the exception of two codon variants for leucine, the presence of two SNPs within the same codon will always lead to a non-synonymous codon. Thus, without codon-constrained SNP linkages, we hypothesized that we may be underestimating the number of genes detected under positive selection using pN/pS. MetaPop tries to resolve this issue by linking SNPs at the read level (as many tools do), but also at the codon level (see methods in Step 3: Microdiversity Analyses section above). We tested our hypothesis on the GOV2 and autism biological datasets using MetaPop and a global SNP calling approach to maximize the number of codons with putatively linked SNPs per sample.

Of the total genes in the autism dataset, ~1.4% of genes (n=248) with enough coverage to evaluate selection had ≥1 codon with putatively linked SNP in at least one sample (**Fig. 3A, top – larger circle**). Of this subset, 16.97% contained at least one codon with potentially linked SNPs (**Fig. 3A, top – smaller circle**). The subset of genes containing a putatively linked codon had their pN/pS ratios calculated using both with and without linking SNPs, and their results were compared. When SNPs were not linked, we observed that 21.7% of the genes (n=54) displayed positive selection, and when linking SNPs, we observed that 26.9% of the genes (n=63) displayed positive selection (**Fig. 3A, bottom**). There were minimal differences using a PHRED≥30 or ≥20, with PHRED≥30 detecting 25.4% (n = 258) genes under positive selection using the codon-constrained SNP linkages, compared with 26.9% (n = 271) with PHRED≥20 (**Fig. 3B**). In GOV2, we observed a similar pattern. In GOV2, 5% of genes (n = 229,058) with enough coverage to evaluate selection had ≥1 codon with putatively linked SNP (**Fig. 3C, top – larger circle**). Of this subset, 20.1% contained at least one codon with potentially linked SNPs (**Fig. 3C, top – smaller circle**). Similarly to the autism dataset, we saw that SNPs linkages increased the number genes detected under positive selection from 21.7% (n=50,094) to 24.56% (n=56,467) (**Fig. 3, bottom**). Thus, MetaPop’s codon-constrained SNP linkage shows that we are underestimating the number of positively selected genes that contain these putatively linked SNPs by an average of ~4%. Further, it shows that utilizing pN/pS to identify genes under selection without linking SNPs at the codon misses genes under positive selection.

**Figure 3.**
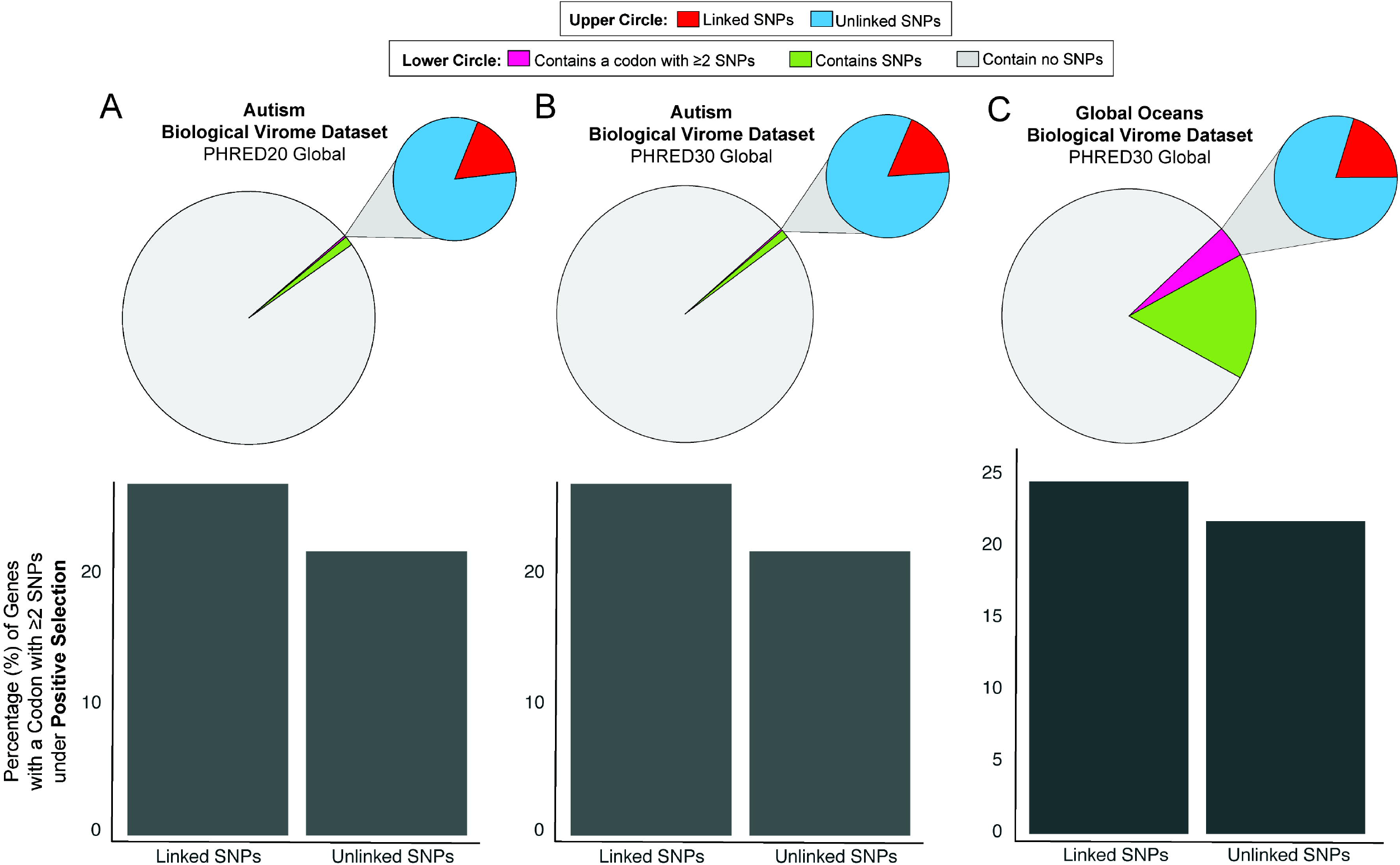
MetaPop’s codon-level SNP linkages increase the number of positively selected detected genes using pN/pS. **(A-C, top)** Pie charts displaying the predicted genes belonging to a contig which passed preprocessing coverage and depth cutoffs, genes with at least one SNP observed, and genes with at least one codon with putatively linked SNPs, and pop-out pie chart showing the breakdown of genes with observed SNPs and those containing a codon with putatively linked SNPs. **(A-C, bottom)** Barplots comparing the percent of genes under selection when calculated after linking SNPs vs. not attempting to link SNPs. Data is shown for the autism biological virome dataset using **(A)** PHRED20 and **(B)** PHRED30 global SNP calls and on the global oceans biological virome dataset using **(C)** PHRED30 global SNP calls.

#### Microdiversity: a case study in assessing intra-population variation reveals gut viruses may play a role in dysbiosis of autistic children’s guts

To demonstrate the value of adding estimates of *micro*diversity to a researcher’s toolkit, we used MetaPop to re-analyze the gut viromes of autistic children that underwent fecal microbiota transfer (FMT) and their neurotypical peers. Up to seventy percent of autistic children suffer gastrointestinal problems, so understanding how the gut microbiota differ between autistic and neurotypical children may be important for treating this symptom of autism (Hsiao 2014). Previously, we found bacterial macrodiversity was lower in autistic children compared to their neurotypical peers, but that viral *macro*diversity (Shannon’s H) was not significantly different (see **Fig. 4A;** Wilcoxon test *p* = 0.89; Kang *et al*. 2017). Thus for this demonstration, we chose to specifically focus on the *macro*-α-diversity Shannon’s H (as a positive control for whether MetaPop could recover our past observations), and the *micro*diversity average π (to assess what biological inferences can be gained by such measures).

**Figure 4.**
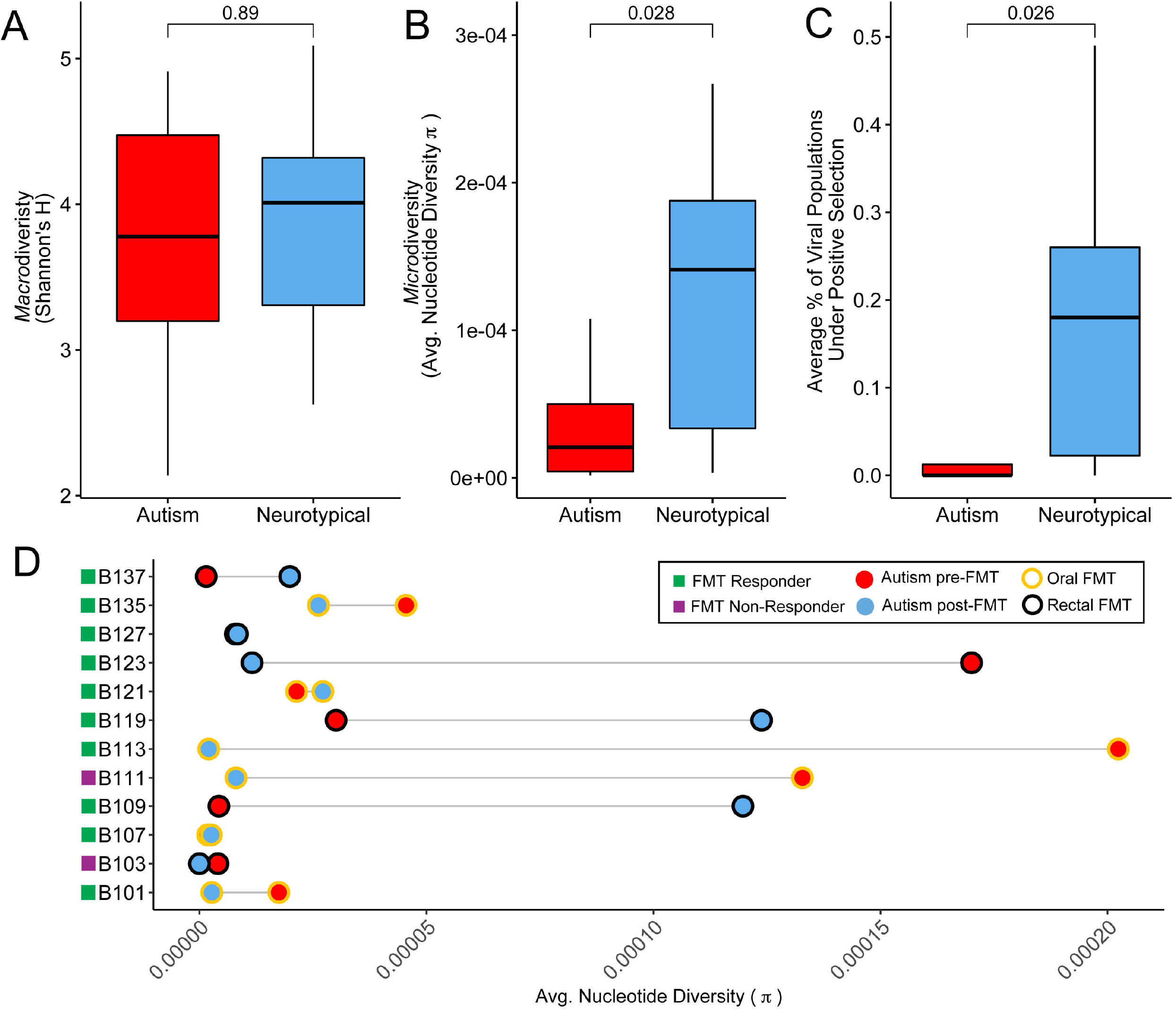
MetaPop’s *microdiversity* analyses reveal lower *intra*-population diversity in autistic children. Boxplots showing median and quartiles of **(A)** Shannon’s H, **(B)** average nucleotide diversity (π), and **(C)** the average number of genomes under selection in pre-FMT autistic and neurotypical children. The Wilcoxon test *p*-values above are the result from comparing pre-FMT autistic and neurotypical children values. **(D)** Connected dot plot showing the average π for the autistic children pre-FMT and post-FMT. The color filling the circle represents the pre- and post-FMT status and the outlier color of the circle represents the method in which the FMT was delivered (oral or rectal delivery). The colored squares next to autistic child’s ID corresponds to whether the child responded or did not respond to the FMT treatment.

Here, MetaPop revealed that average viral *micro*diversity (π) is significantly lower in autistic children than within their neurotypical peers **(Fig 4B;** Wilcoxon test *p* = 0.028), paralleling our previous bacterial *macro*diversity findings (Kang *et al*. 2017). High average π can indicate two biological outcomes: (*i*) more viruses from different populations are actively infecting host bacteria resulting in population expansion and more mutations, or (*ii*) more viral populations naturally maintain higher levels of *micro*diveristy in their standing populations. Either way, having a higher level of *micro*diveristy for a viral population can be an adaptive advantage because it better allows populations to ‘bet hedge’ if their environment or hosts change (Hughes *et al*. 2008). We next looked to see if this increased *micro*diversity could be providing an adaptive advantage to the neurotypical children by exploring the average number of genomes containing a gene found under positive selection using pN/pS. Indeed, neurotypical children had significantly more genomes with at least one gene under selection than the autistic children **(Fig 4C;** Wilcoxon test *p* = 0.026). Thus, we hypothesize that increased average π is beneficial in the gut virome because it allows viral populations to better adapt to their changing environments and hosts. Increased diversity at either the *macro*- and *micro*-diversity level has consistently been shown to be important for maintaining ecosystem functions and services (e.g. Tilman *et al*. 2014, Hughes *et al*. 2008), with the loss of diversity resulting in a loss of ecosystem resilience. Thus, the loss of viral *micro*diversity in autistic children’s guts could potentially be indicative of a loss of the gut ecosystem’s resilience.

Next, given that the autistic children underwent FMT, we were also curious how FMT impacted their viral *micro*diversity. We compared the pre-FMT (week 0) to the post-FMT (week 10) gut viral *micro*diversity (see Kang *et al*. 2017 for full information about FMT design). Of the 12 autistic children with viromes available, 10 of the children responded positively to FMT treatment and 2 did not. Given the hypothesis that increased average π is beneficial, we expected to see that all of the responders would have increased viral *micro*diversity. Instead, we saw an interesting pattern (**Fig 4D)**. Six of the children with virome data were given the FMT orally, with the other six were given the FMT rectally. Across the responders that received FMT orally, only 2 of the 5 children had increased viral *micro*diversity, and only an average 1.36-fold increase at that. In contrast, 4 of the 5 children that received the FMT rectally responded with increased viral *micro*diversity, and they did so with a larger (11.58-fold) average increase. This suggests that rectal administration of the FMT may promote engraftment of more *micro*diverse viral populations, at least those surveyed in the feces, than the oral administration of FMT. This contrasts the findings at the clinical symptom-level that found no significant difference in changes in children that received the oral or rectal FMT (Kang *et al*. 2017). Taken together, though the study was pilot-scale and open-label, these *micro*diversity results suggest that viral population structure is associated with the autism disease phenotype and that FMT delivery method may correlate to responder status. Again, however, a larger study is needed to better guide standard of care practices.

### Computational Evaluation of MetaPop

We next evaluated the processing time and computational resource consumption of MetaPop by running the synthetic mock bacterial and the biological virome datasets on the Ohio Supercomputer (OSC). To simulate different computational power, from a standard laptop to desktop computer to a small sized computer cluster, we ran MetaPop using 4, 6, and 14 cores, using 8, 16, and 64 GB of memory (RAM), respectively (**see Fig. 5A**). Notably, the much larger GOV2 dataset needed more cores and memory to run than the different settings tested (~1.5TB), so it was excluded from this analysis. Importantly, MetaPop’s code is parallelized, so increasing the number of supplied cores will increase the memory used because MetaPop will try to parallelize its steps as efficiently as possible given the computational resources supplied.

**Figure 5.**
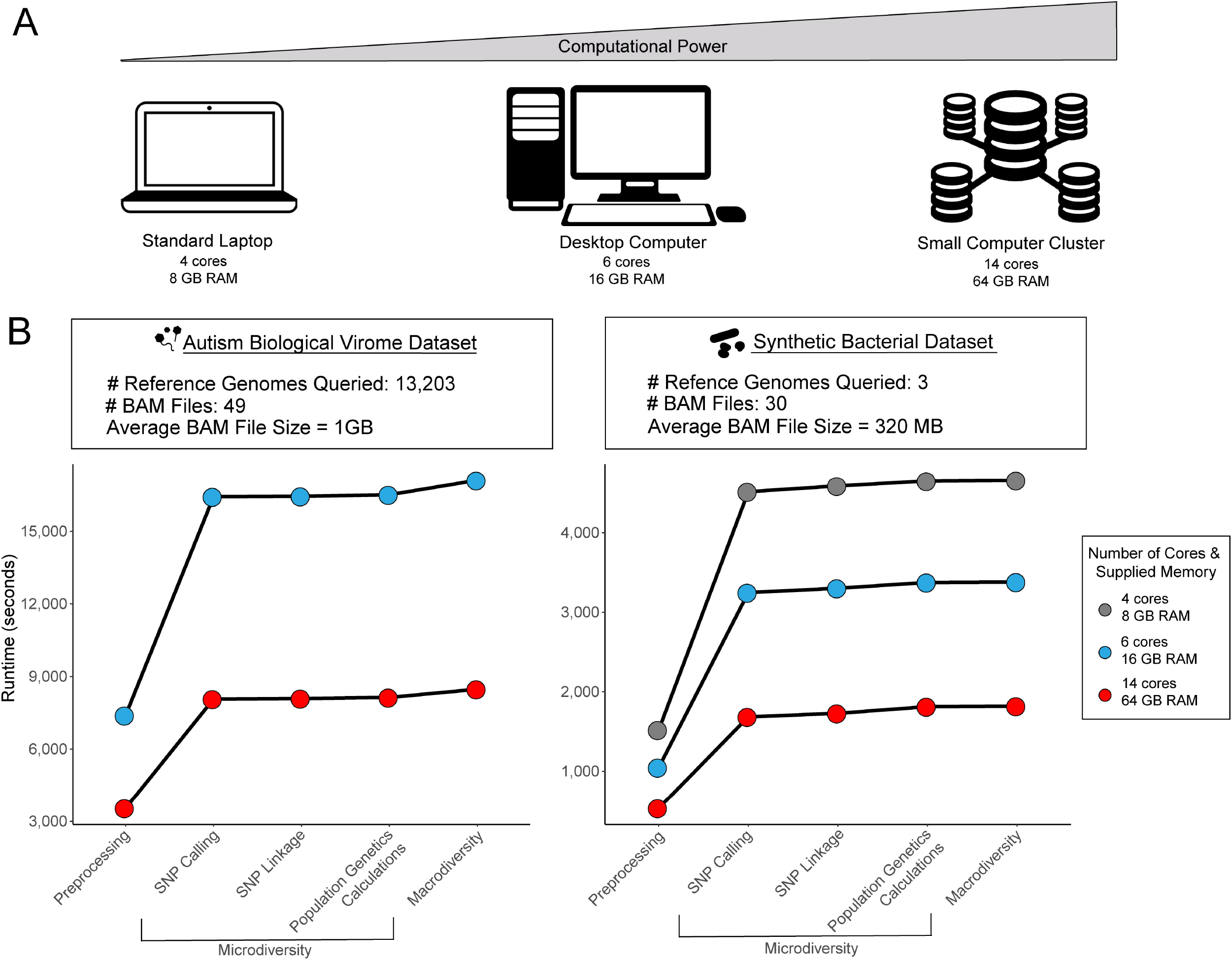
Computational evaluation of MetaPop. **(A)** Schematogram showing the different computational power levels tested. **(B)** Line plots showing cumulative runtime of MetaPop on **(left)** the biological dataset with 6 and 14 processors and **(right)** the synthetic dataset with 4, 6, and 14 processors.

To assess both processing time and computational resource consumption, we first wanted to determine what steps in the MetaPop pipeline were rate-limiting. Across both the synthetic and biological datasets, the pre-processing and the SNP calling portion of the microdiversity section took the most processing time (**Fig. 5B – right and left**). These two steps operate on the entirety of data supplied by the user, and must perform multiple operations on every read in each BAM file, and for every contig supplied. This means that they must process large volumes of data and consume commensurate computational resources. Further, the resources consumed by these steps depend on the degree of parallelization, as each parallel process will operate on its own set of data, simultaneously. The latter portions of the pipeline work on summaries produced by pre-processing and SNP calling steps and are, correspondingly, faster because they do not have to operate on the entire input dataset, even though these later steps have substantially less parallelization.

With the knowledge of the rate-limiting steps, we then assessed the effect of BAM file size and the amount of computational power supplied on processing times. BAM files size is impacted by the number and length of reference genomes and the original number of reads. The synthetic and biological datasets differ substantially in these values resulting in vastly different BAM file sizes and allowing us to test MetaPop across a range of BAM file sizes, with the average synthetic dataset’s BAM file size equal to 320 megabytes (MB) and the average biological dataset’s BAM file size equal to 1 gigabyte (GB). The processing time of the rate-limiting steps were nearly linear functions of BAM file size across the biological dataset (linear regression: R^2^ =0.902; **Fig. S4A**) and the synthetic dataset (linear regression: R^2^=0.796; **Fig. S4B**).

We were next curious about how many of the computational resources were consumed of the memory supplied. The maximum memory consumption for the synthetic dataset was 4.6 GB when run with 14 cores. For the biological dataset, MetaPop failed using the standard laptop computational power because running the pipeline exceeded the 8GB of memory supplied. Using 6 cores and 16GB memory supplied, MetaPop efficiently parallelized and used all 15.05 GB memory. At the higher end of computational power (14 cores and 64GB memory supplied), it is clear that MetaPop did not need all the memory supplied to run efficiently and only used 23.66GB total. Overall, these results show that MetaPop can be successfully run using low computational resources and will adjust the resources consumed based on the computational power supplied for datasets with average BAM sizes around 1GB.

## Limitations and future directions

While MetaPop provides the sort of ease-of-use and scalability that we hope will open up microdiversity analyses to more researchers, our current implementation will benefit from future improvements. *First,* MetaPop was designed and optimized for single-contig genomes, so it does not work with binned-contig genomes and will treat each new contig as a different population. Nonetheless, it is possible to derive *macro*- and *micro*-diversity calculations across binned contigs. The average read depth and nucleotide diversity per position for each contig per BAM file is output, so it is possible to derive the *macro*diveristy abundance tables and *micro*diversity π values for binned contigs from MetaPop output. *Second,* MetaPop’s default settings are optimized for short-read datasets. As more hybrid sequencing and assembly efforts are used to explore microbial and viral communities (e.g., Warwick-Dugdale *et al*. 2019, Moss *et al*. 2020), which capture more niche-defining hypervariable regions, adjustments for MetaPop’s abundance calculations and SNP calling will need to be done. Though not prohibitive, this will require benchmarking studies that assess the nuances of new sequencing technology (e.g., homopolymers for nanopore sequencing) to correct for per-base-pair sequencing errors against the background of real biological mutations, many of which are now emerging (e.g., Amarasinghe *et al*. 2020). *Finally*, MetaPop is benchmarked for studying community and population-level diversity and selection, we have not optimized it for resolving strain-level genotypes. However, as other tools (e.g. InStrain; Olm *et al*. 2020) solve the problem of reconstructing strains from metagenomes, the resultant genotypes could be input to MetaPop.

## Conclusions

Metapop is a fast and scalable pipeline for the analyses of both *macro*- and *micro*diversity in metagenomic data. It combines both classical community ecology metrics with the full suite of population genetics parameters in a single integrated pipeline. While many of its functions are already available in existing pipelines (Eren *et al*. 2015, Costea *et al*. 2017, Nayfach *et al*. 2016, Olm *et al*. 2020), MetaPop’s easy user-interface (i.e. single-line command) and ability to be run on a standard laptop for smaller datasets make it a practical choice for non-bioinformaticians and microbiology labs without access to large supercomputers. Further, MetaPop’s default visualizations enable fast and easy interpretation of the results.

Molecular biology and sequencing technology advances have advanced the microbiologist’s toolkit from 16S rRNA gene analyses to metagenomics and changed questions we could ask from “who is there” to also add “what could they be doing” and “with whom might they interact”. Now, with further technological advances and by democratizing microdiversity analyses, we open a new window into the study of complex communities such that we can now ascertain “what populations have high *micro*diversity levels” and “which genes are under selection”. While studying *micro*diversity has been hard due to lack of data and ease of toolkit, we are entering an era where such data and toolkits are available such that our understanding of these new biological windows will provide new insights into how complex systems work. MetaPop gives scientists an ideal toolkit to explore the dual impact of *macro*- and *micro*diversity across microbially-impacted ecosystems.

## Materials and Methods

### Preparing the mock and biological dataset input files for MetaPop

We chose three previously published datasets, a synthetic dataset representing mock bacterial communities (Sankar et al. 2016) and two biological virome datasets including 131 viromes from the global oceans virome 2 (GOV2) datasets (Gregory *et al*. 2019) and gut viromes from autistic children that underwent FMT and their neurotypical peers (Kang et al. 2017), to evaluate MetaPop. The synthetic dataset was composed of known proportions of three distinct strains of *S. aureus* (ST5, ST8, and ST30), three distinct strains of *S. epidermidis* (TAW60, CV28, 1290N), and a single strain of *B. subtilis* (Sankar *et al*. 2016). Because MetPop explores inter- and intra-population-level analyses, we selected one strain of *S. aureus* (ST5 strain ECT-R2; GenBank: NC_017343.1), one strain of *S. epidermidis* (TAW60: binned assembly from Méric *et al*. 2015), and one strain of *B. subtilis* (strain 168; GenBank: NC_000964.3) as the population-level genome representatives. For the GOV2 dataset, we used the *Tara Oceans* 131 viromes from GOV2 and did not process the *Malaspina* viromes. The 488,130 viral populations identified in Gregory *et al*. 2019 were used as the reference genomes. For the biological gut virome dataset which comprised 49 gut viromes, we used the gut viral database from Gregory *et al*. 2019 as the population-level reference genomes. Reads form both the mock and biological datasets were non-deterministically read mapped to their respective reference genomes using bowtie2 (Langmead *et al*. 2012). The resulting BAM files, the reference genomes, and read counts for each metagenome were used as input for MetaPop. MetaPop was run using the default settings if not otherwise noted.

### Evaluating the processing time, computational resource consumption, and scalability of MetaPop

In order to computationally evaluate and benchmark MetaPop, we emulated the resource environment of several likely computational platforms and attempted to process both the synthetic and biological datasets using these resources. The average BAM file size of the mock synthetic dataset BAM is 320 MB and has 30 samples, and of the biological dataset BAM is 1013 MB, with a total of 49 samples. The chosen computational scales reflect a fairly typical laptop computer, with 4 processing cores and 8 GB of RAM, a desktop computer with 6 cores and 16 GB or RAM, and a supercomputing environment using all 14 cores on one of the Ohio Super Owens (OSC) nodes and 64 GB of RAM.

The Owens nodes are each equipped with a Intel Xeon e5 2680 v4 Broadwell processor, which has 14 cores, and each node shares identical RAM and data storage characteristics. This provides parity between the differing scales of the computing environment, rendering maximum permissible memory and allocated cores the sole difference affecting runtime and memory usage. Finally, the OSC process manager terminates the execution of code which exceeds the supplied memory of any given job, meaning that exceeding specified RAM results in a failure of MetaPop to complete, just as it would in an environment actually limited by such resources. While the manager would also terminate a job which exceeded a specified runtime, we supplied each instance with excessive runtime so that this would not be a factor.

During preprocessing, MetaPop creates a tab separated file that records the start and end dates and times of each of the 5 core components of its code, namely the preprocessing, SNP calling/refinement, linked SNP read mining and linkage calculations, calculation of microdiversity, and the combined calculation of macrodiversity and creation of macrodiversity visualizations. This file provides accurate timing for the overall runtimes of each processing phase, and served as the basis for timing the pipeline as a whole. These files were simply combined and organized to produce the overall runtimes for MetaPop at each processing scale and stage.

In order to time the individual runtimes of MetaPop where processes are independent of each other, a modified version of the core MetaPop code was used. The modifications made to the code are simple system calls to print the start and end date and time to a series of text files where individual runtimes are applicable. These calls occur within microseconds, and thus do not substantively affect overall runtimes.

In order to profile memory usage during the various phases of MetaPop, we relied on computational resource logs produced by the Ohio Supercomputer. These files report a variety of computational resource consumption statistics associated with a particular task, which includes the peak memory footprint for any collection of processes contained within a single job on the supercomputer. This approach answers the most pertinent question for users: what is the minimum RAM that is required to run a dataset of a given scale through MetaPop with a particular number of cores.

### Mock Community Macrodiversity Validation

In order to assess MetaPop’s ability to resolve macrodiversity, we compared MetaPop’s predicted relative abundances to the known relative abundances in the 30 mock communities (Sankar *et al*. 2016). Importantly, some of the reference genomes for the mock communities were not closed genomes. Thus, in order to calculate the raw abundances, the mean coverage across all base pairs in all fragments within the reference gene were calculated, excluding coverages below the 10th and above the 90th percentile per base pair. These values were then scaled as described in the ***Macrodiversity analyses*** above to create the normalized abundances. Prior to comparing the relative abundances, MetaPop’s calculated normalized abundances were converted into relative abundances by dividing each population’s normalized abundance in a community by the sum of all the population’s normalized abundances. The fold-change difference between MetaPop’s calculated observed relative abundances and known relative abundances was then assessed using ‘foldchange’ in the R package ‘gtools.’ Next, the calculated *macro*diversity α-(Richness, Shannon’s *H,* and Peilou’s J) between the observed MetaPop values and the expected actual values across the 30 mock communities were compared using Wilcoxon tests in the R package ‘ggpubr.’ β-(Bray-Curtis dissimilarity) diversity calculated distances calculated across all 30 communities then compared using a Mantel’s test in R package ‘vegan.’ Lastly, FastANI (Jain *et al*. 2018) using default settings was used to compare average nucleotide identity across the different strains and species within the mock community.

### Mock Community Codon Bias Analyses

We chose to evaluate MetaPop’s ability to pull out genes with different codon bias usage by examining the *Staphylococcus aureus* strains in the 30 mock communities (Sankar *et al*. 2016). *S. aureus* are well studied to their clinical relevance and there is a great deal of knowledge about the different genes and elements that have been horizontally acquired within their genomes (Malachowa *et al*. 2010; Alibayov *et al*. 2014) and, to some extent, genes with increased expression (Malachow *et al*. 2011, Karlin *et al*. 2001; Soufo *et al*. 2010). In order to assess the codon bias usage outlier part of MetaPop, we manually curated a list of known horizontally transferred and highly expressed genes and compared it to the list of genes with different codon bias usage predicted by MetaPop. We then calculated what proportion of the predicted genes with different codon bias usage were known to be horizontally acquired or highly expressed.

### Global Oceans Viromes 2 (GOV2) Microdiversity Validation

In order to assess MetaPop’s ability to resolve *micro*diversity values and patterns, we computed the *micro*diveristy (average π) per sample in the 131 GOV2 samples based on MetaPop’s calculated nucleotide diversity (π) and compared it to the published *micro*diversity values (Gregory *et al*. 2019). To calculate *micro*diveristy for each sample, average π was calculated by randomly selecting the π values of 100 viral populations and then averaging their values (*sensu* Gregory & Zayed *et al*. 2019). This was repeated 1000x and the average of the all 1000 subsamplings was used as the final average *micro*diversity value for each sample. The values were plotted using the line graph and scatter plot functions in Excel. The linear regressions were also run in Excel. The sample *micro*diversity values were then grouped by ecological zone as defined in Gregory *et al*. 2019 and, unlike the original analyses, the values were not subsampled from each ecological zone and then averaged in order to see the better spread of the values per zone. The ecological zone values were plotted and statistical differences assessed using the R package ‘ggboxplot.’ GOV2 SNPs were originally locally called using a PHRED ≥ 30. We ran MetaPop using a PHRED ≥ 20 (MetaPop’s default) and ≥30 to filter the variants and then assessed SNP calls both globally (for PHRED≥30 and ≥20) and locally (for PHRED≥30). The fold-change difference between MetaPop’s average π values and the original GOV average π values were then assessed using ‘foldchange’ in the R package ‘gtools.’

### Biological Dataset Microdiversity Analyses

To compare macro- and micro-diversity in the autism virome dataset (Kang *et al*. 2017), the predicted *macro*diversity α- (Shannon’s H) values, the *micro*diveristy (average π), and the percentage of genomes under positive selection for all the autistic and neurotypical children prior to FMT treatment were compared using Wilcoxon tests in the R package ‘ggpubr.’ To calculate *micro*diveristy for each sample, average π was calculated by randomly selecting the π values of 10 viral populations and then averaging their values *(sensu* Gregory & Zayed *et al*. 2019). This was repeated 50x and the average of the all 50 subsamplings was used as the final average *micro*diversity value for each sample. To calculate the percentage of genomes under positive selection per sample, viral populations with at least one gene detected under positive selection (pN/pS > 1) per sample were determined and pooled with the viral populations with enough coverage to analyze *micro*diversity. Similarly to average π, the 10 viral populations per sample were randomly selected and then the percentage that were detected to be under positive selection assessed. This was repeated 50x and the average of the all 50 subsamplings was used as the final average percentage of viral populations detected under positive selection for each sample. The pre- and post-FMT viromes of the autistic children were then plotted using the R package ‘ggplot2’ and the fold-change difference was assessed using ‘foldchange’ in the R package ‘gtools.’

## Data Availability

MetaPop is available for download at https://github.com/metaGmetapop/metapop and can be downloaded as the bioconda package ‘metapop’. The synthetic datasets used for this study are available at: http://figshare.com/articles/Benchmarking_data_for_bacterial_strain_identification/1617539.

The 131 GOV2 viromes used in this study can be downloaded from the European Nucleotide Archive (ENA) under the accession numbers found in Supplementary Table 3 of Gregory *et al*. 2019. The virome datasets used for this study are available in iVirus at the following link: http://mirrors.iplantcollaborative.org/browse/iplant/home/shared/iVirus/ABOR. Support for the pipeline is available on the issue tab of the github page.

## Supplementary Data

Supplementary Data is available online.

## Acknowledgments

Pipeline design and discussion with Sergei Solonenko and Cesar J. Ignacio Espinoza is gratefully acknowledged. For help with digging into the *Staphylococcus* literature, we would like to thank Rodrigo Bacigalupe and Amy Richards. Lastly, we would like to thank the Sullivan lab and Konstantinidis lab members for testing the pipeline and providing input during the development of MetaPop.

## Main Text Tables

See Excel document for table (https://drive.google.com/file/d/1okwESg4Qf5mq0oaB0Ud-gFRPgAKfKpaL/view?usp=sharing)

**Table 1.** Capabilities of MetaPop compared to existing complementary bioinformatic pipelines.

