## Supplemental Figures and Tables for "MetaPop: A pipeline for *macro*- and *micro*-diversity analyses and visualization of microbial and viral metagenome-derived populations"

**Supplementary Figures:**

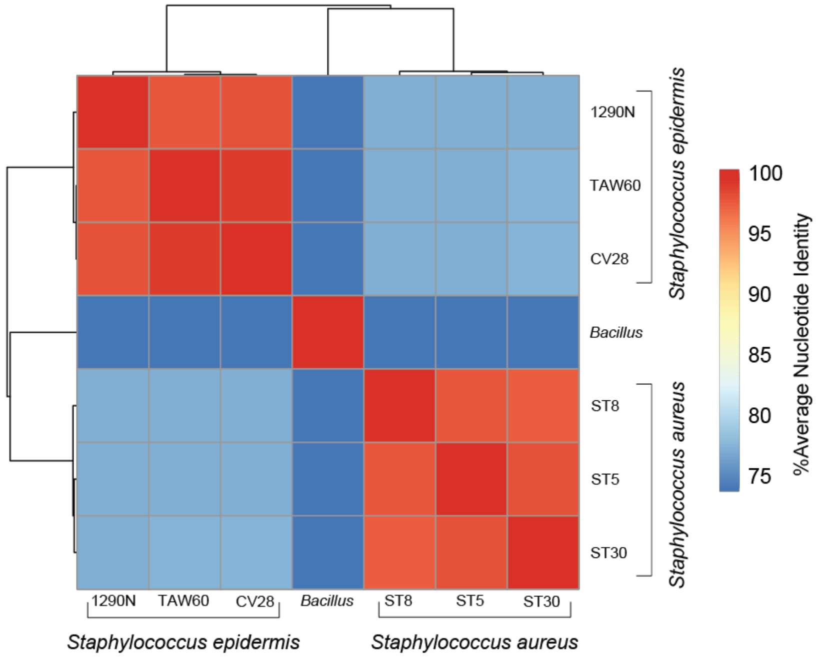


**Figure S1.** Heatmap showing % average nucleotide identities (ANI) similarities among the different strains and populations in the 30 mock communities

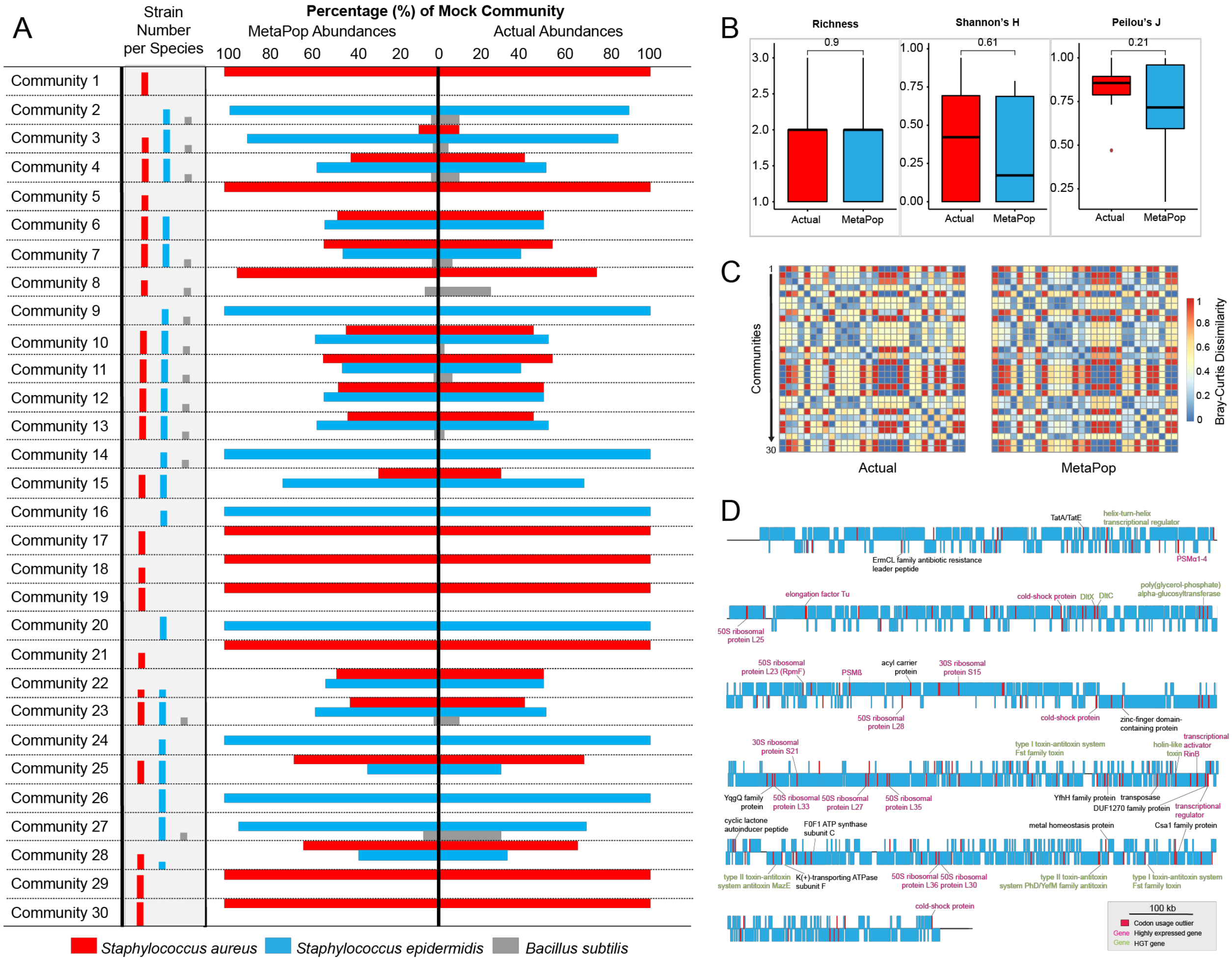


**Figure S2. Validating MetaPop's *macro*diversity and codon bias analyses. (A)** Tornado plot showing the relative abundances of *Staphylococcus aureus*, *Staphylococcus epidermidis*, and *Bacillus subtilis* across the 30 mock communities in the actual synthesized community and as determined by MetaPop. Bar charts contained within the gray bar to the left of the tornado plot reveal the number strains per each bacterial species, with three being the highest number of strains per species. **(B)** Boxplots showing median and quartiles of different α-diversity indices (richness, Shannon's H, and Peilou's J) compared between the actual and MetaPop derived abundances. The Wilcoxon test *p*-values above are the result from comparing actual and MetaPop derived α-diversity indices. **(C)** Heatmaps of β-diversity Bray-Curtis dissimilarity distances calculated using the actual and MetaPop derived abundances. **(D)** Genome map of genes with outlier codon usage in ST5 *Staphylococcus aureus* ECT-R2.

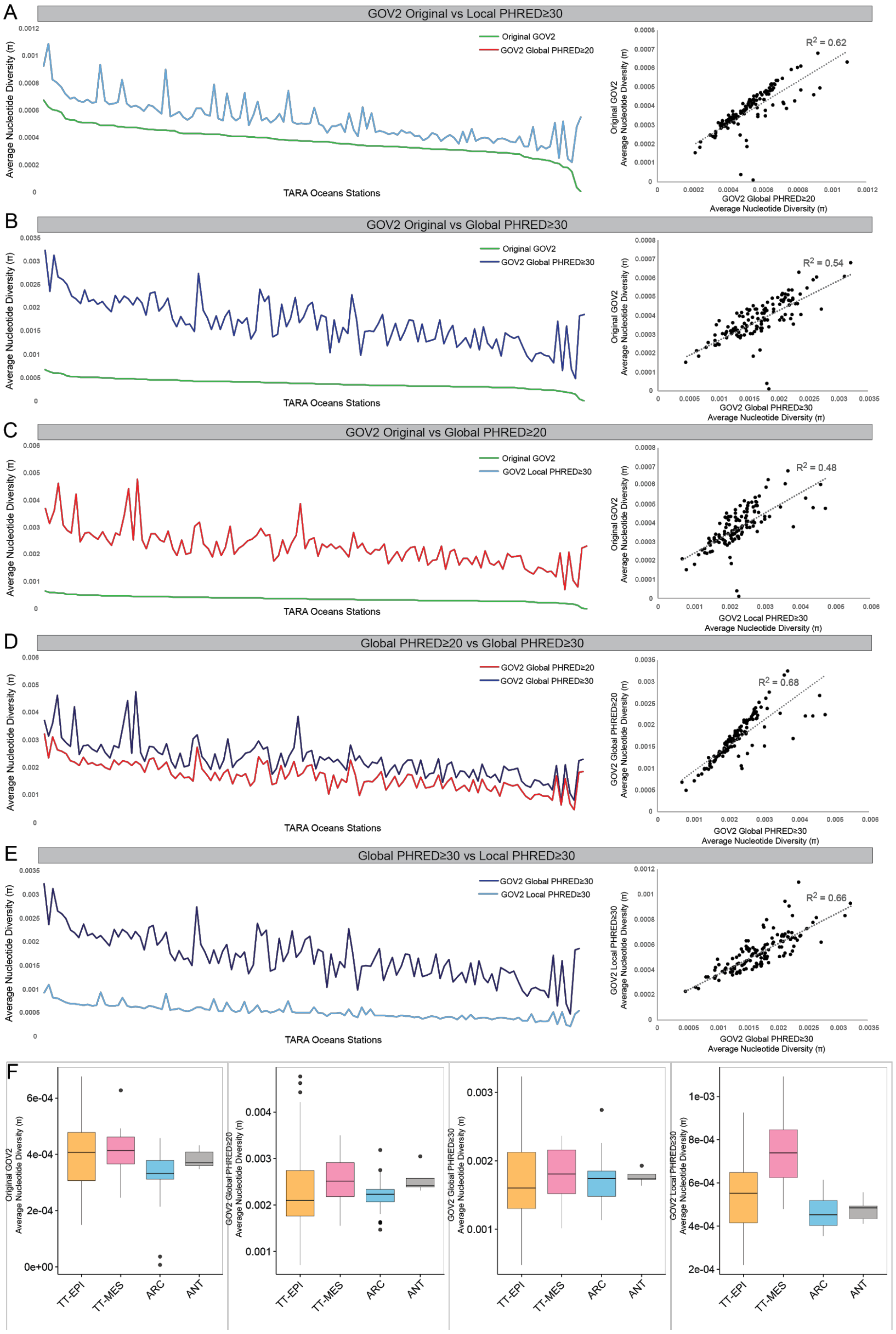


**Figure S3. Validating MetaPops's microdiversity analyses using the Global Oceans Virome 2 dataset. (A-E, right)** Line plots sorted by the original average nucleotide diversity (π) values from Gregory *et al.* 2019 and **(A-E, left)** scatter plots comparing the average π for the *Tara* Oceans stations in the GOV2 dataset derived from the **(A)** original GOV2 values versus MetaPops's PHRED≥30 local SNP calls, **(B)** original GOV2 values versus MetaPops's PHRED≥30 global SNP calls, **(C)** original GOV2 values versus MetaPops's PHRED≥20 global SNP calls, **(D)** MetaPops's PHRED≥20 global SNP calls versus MetaPops's PHRED≥30 global SNP calls, and **(E)** MetaPops's PHRED≥30 global SNP calls versus MetaPops's PHRED≥30 local SNP calls. The dashed line in the scatter plot represents the linear regression. **(F, left to right)** Bar plots showing the biological *micro*diveristy trends across the ecological zones defined in Gregory *et al.* 2019 derived from the original GOV2 values, MetaPops's PHRED≥20 global SNP calls, MetaPops's PHRED≥30 global SNP calls, and PHRED≥30 local SNP calls.

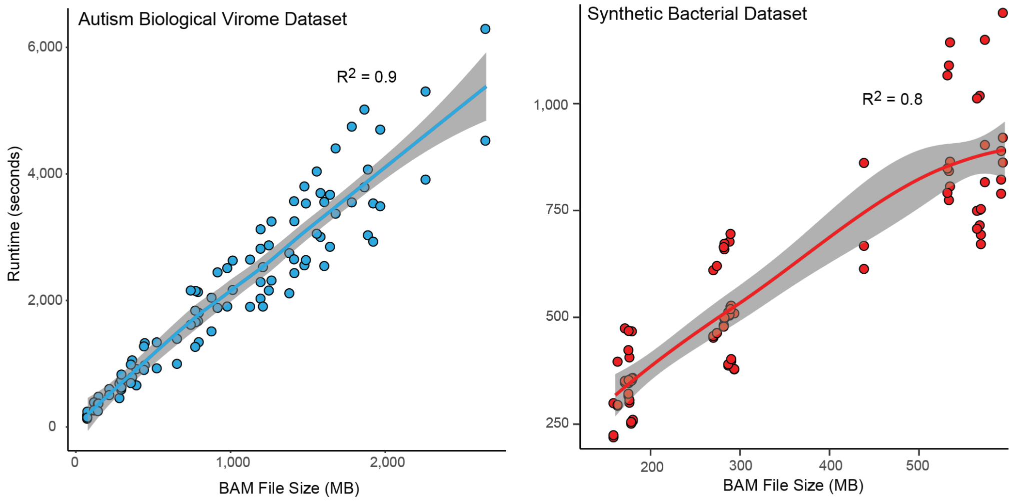


**Figure S4.** Scatterplots with Loess smoothing displaying runtime per sample for the rate-limiting part of MetaPop (i.e. pre-processing and the SNP calling section of microdiversity) as a factor of file size in megabytes on **(left)** the biological dataset and **(right)** the synthetic dataset.

**Supplementary Tables:**

See Excel document for table (https://drive.google.com/file/d/1okwESg4Qf5mq0oaB0Ud-gFRPgAKfKpaL/view?usp=sharing)

**Table S1.** Mock communities data actual and metapop derived abundances.
**Table S2.** Biological virome dataset population abundances.
**Table S3.** Full codon bias usage results for all genes in *Staphylococcus aureus* ECT-R2.
**Table S4.** MetaPop processing times with different levels of computational power.
